# Single-Molecule Manipulation of Macromolecules on Membranes Using High-Resolution Optical Tweezers

**DOI:** 10.1101/2021.07.15.452421

**Authors:** Yukun Wang, Huaizhou Jin, Yongli Zhang

## Abstract

Despite their wide applications into soluble macromolecules, optical tweezers have rarely been used to characterize dynamics of membrane proteins, mainly due to lack of model membranes compatible with optical trapping. Here, we found that optical tweezers can stably trap giant unilamellar vesicles (GUVs) containing iodixanol with controlled membrane tension, which can potentially serve as a model membrane to study dynamics of membranes, membrane proteins, or their interactions. We also observed that small unilamellar vesicles (SUVs) are rigid enough to resist large pulling force and offer potential advantages to pull membrane proteins. To demonstrate the use of both model membranes, we pulled membrane tethers from the trapped GUVs and measured the folding or binding dynamics of a single DNA hairpin or synaptotagmin-1 C2 domain attached to the GUV or SUV with high spatiotemporal resolution. Our methodologies facilitate single-molecule manipulation studies of membranes or membrane proteins using optical tweezers.

## INTRODUCTION

Numerous biological processes on membranes involve complex protein-protein and protein-membrane interactions that are further regulated by mechanical force. These processes include membrane protein folding (1, 2), membrane fusion or lipid exchange (3–5), cell migration (6), immune responses (7), mechanosensation (8–10), and mechanotransduction (1, 5, 8–11). These interactions are difficult to study using traditional experimental approaches based on a large number molecules due to ensemble averaging or lack of mechanical force (3–5, 12–14). Single-molecule force spectroscopy, including atomic force microscopy (AFM), optical tweezers, and magnetic tweezers, has widely been applied to study dynamics of soluble proteins (15–22).

However, the methodology has much less applications into membrane proteins. AFM can image membranes, apply force to membrane proteins, and probe protein dynamics (23). Consequently, AFM has long been used to study membrane protein folding by pulling single proteins out of surface-supported lipid bilayers (24, 25). Yet, reversible protein folding into membranes has rarely been observed, except for small regions of transmembrane helices, which prevents measurements of folding energy for larger domains of membrane proteins, including the insertion energy of a single transmembrane helix. AFM generally uses large and stiff fabricated cantilevers as force probes, which lead to high spatial resolution, but low force resolution compared with magnetic or optical tweezers (17, 26). In addition, the supported bilayers may introduce artifacts due to protein-surface interactions, which may perturb the structure and dynamics of embedded membrane proteins, including reduced lateral diffusion (27–29). Magnetic tweezers have been successfully applied to detect stepwise association and dissociation of transmembrane helices of rhomboid protease GlpG or β2-adrenergic receptor in bicelles, and recently, unfolding of GlpG in small unilamellar vesicles (SUVs) (30, 31). Yet, dynamic protein unfolding out and folding into membranes has not been observed. Compared with AFM and magnetic tweezers, optical tweezers are most widely used to study dynamics of soluble proteins, including unidirectional movement of molecular motors and folding dynamics of proteins or protein complexes (16, 21, 22), partly due to their extremely high precision for measurements of distance (~0.2 nm) and force (~0.01 pN) with extraordinary temporal resolution (~10 microseconds) (32). In contrast, optical tweezers are also least used to investigate membrane proteins, especially their folding dynamics, partly due to lack of proper model membranes to be suspended in optical traps to pull membrane proteins. Our objective was to develop such model membranes to facilitate studies of membrane proteins using optical tweezers.

Giant unilamellar vesicles (GUVs) and SUVs are common model membranes to study membrane proteins in bulk (33). Integral proteins on both GUV and SUV membranes are freely mobile (34). Aspirated on the tips of micropipettes, GUVs have been utilized as membrane reservoirs to pull long membrane tethers or tubules with controllable diameters or curvatures with optical tweezers (35–37). These membrane tethers serve as ideal substrates to test many proteins that bind to membranes in a curvature-dependent manner or deform the membranes upon their binding (38). Micron-sized GUVs could also be trapped by optical tweezers to probe the mechanical properties of lipid bilayers (39). However, the optical trapping was weak, due to the small difference in the refractive indices (RI) of GUVs and water, making it unlikely to directly pull membrane proteins reconstituted onto the trapped GUVs. Furthermore, reconstitution of integral membrane proteins into GUV membranes is generally challenging, as there has been no general methods for reliable protein reconstitution (40). SUVs are popular model membranes for membrane protein studies, partly because reconstitution of membrane proteins onto SUVs is generally easier. However, with a diameter ranging from 20 to 100 nm, SUVs are invisible by conventional optical microscopy and difficult to be trapped by optical tweezers. Taken together, it remains challenging to pull single membrane proteins using optical tweezers.

We developed methods to pull macromolecules attached to the membranes of GUVs and SUVs to measure dynamics of proteins and/or membranes using optical tweezers with high resolution. We validated our methods using DNA hairpins and synaptotagmin-1 and demonstrated the potential applications of both model membranes to studies of integral membrane proteins using optical tweezers.

## MATERIALS AND METHODS

### DNA handles

Four DNA handles used in the different experiments had the same length of 2,260 bp and dual digoxigenin labels at one end, but different overhang oligonucleotides and/or labels (biotin or thiol group) at the other end. All DNA handles were made by polymerase chain reaction (PCR) using λ DNA cl857 Sam7 (Promega, D1501) as a template and a forward primer containing two digoxigenin labels at the 5’ end. Four reverse primers contained either the overhangs and/or biotin or thiol labels at the 5’ end. The DNA handle used in Fig. 2A had an overhang DNA hairpin sequence of biotin-5’-TTTGAGTCAA-CGTCTGGATC-CTGTTTTCAG-GATCCAGACG-TTGACTCTTT-(spacer), while the left DNA handle in Fig. 5A contained an overhang sequence 5’-CTCGCCAACG-TACATACAAC-TGTACGCCCTC-(spacer). Here the 18-atom hexa-ethyleneglycol spacer connected the overhangs to the DNA handles but prevented polymerase extension to the overhang regions during PCR. All primers were synthesized in IDT (Integrated DNA Technologies, Inc.).

### DNA oligo-lipid conjugation

The DNA hairpin oligo was labelled to lipid (Fig. 5A) via the maleimide–thiol reaction (41). The oligonucleotide with a 3’ thiol group was synthesized by IDT with the following sequence: 5’-GAGGGCGTAC-AGTTGTATGT-ACGTTGGCGA-GTTGAGTCAA-CGTCTGGATC-CTGTTTTCAG-GATCCAGACG-TTGACTCT-SH. The lyophilized oligonucleotide was dissolved in the buffer containing 20 mM Tris, pH7.4, 250 mM KCl, and 55 mM glucose (Buffer A) plus 20 mM TCEP for a 4 mM stock solution. The maleimide labeled lipid 1,2-dioleoyl-sn-glycero-3-phosphoethanolamine-N-[4-(p-maleimidophenyl) butyramide (MPB-PE) dissolved in chloroform in a clear glass vial was dried first in nitrogen flow for 5 minutes and then in a vacuum desiccator for 1 h. All lipids were purchased from Avanti Polar Lipids, Inc. Before lipid labelling, the stock oligonucleotide solution was diluted to 0.8 mM with Buffer A plus 2.5% w/v n-Octyl-β-D-Glucoside (OG) and added to the glass vial covered by the dried lipid film with a MPB-PE:oligonucleotide molar ratio of 10:1. The solution was gently vortexed at room temperature for 4 hours to complete the maleimide–thiol reaction. Finally, the reducing agent 2-Mercaptoethanol was added to the mixture to a final concentration of 40 mM to quench all the unreacted MPE-PE. The oligonucleotide labeled PE was aliquoted and stored in −80 °C before use.

### SUV preparation

SUVs were made for direct use (Fig. 5A) or preparation of MCBs (Fig. 2D) or VAMP2-anchored GUVs or MCBs (Fig. 1C). Three types of SUVs were prepared containing either pure lipids (Fig. 2C), oligonucleotide-PE (Fig. 5A), or VAMP2. In particular, the oligonucleotide-labeled SUVs (Fig. 5A) contained 98.47 mol% 1-palmitoyl-2-oleoyl-sn-glycero-3-phosphocholine (POPC), 1 mol% the oligonucleotide-PE, 0.5 mol% 1,2-dioleoyl-sn-glycero-3-phosphoethanolamine-N-(lissamine-rhodamine-B-sulfonyl) (Rhodamine-PE), and 0.03 mol% biotin-PE. To make all SUVs, different lipids (expect for oligonucleotide-PE) were mixed in chloroform and dried to form lipid films as described in the preceding section. Then Buffer A was added to hydrate the lipids to make a solution with a total lipid concentration of 5 mg/ml. The cloudy vesicle solution was sonicated with a water bath sonicator for 30 min until the solution became clear. These SUVs were ready for use. For SUVs containing oligonucleotide-PE, Triton X-100 (Thermo Scientific, 28314) was added to the SUV solution to a final concentration of 8 mM and incubated for 10 minutes at room temperature with gentle agitation. Then oligonucleotdie-PE was added to the SUV solution to 1 mol% total lipid concentration and further incubated at room temperature for 1 hour. Triton X-100 was removed by adding 40 mg Bio-beads (Bio-Rad Laboratories, 1523920) per 100 μL SUV solution and then nutating at 4 °C overnight. VAMP2-anchored SUVs were prepared as previously described (42). Briefly, the purified Alexa Fluor 647 labeled VAMP2 in 1.5% (w/v) OG, 140 mM KCl, and 25 mM HEPES, pH 7.4 was added to the dried lipids for a total lipid concentration of 5 mg/ml and a protein-to-lipid molar ratio of 1:1000. The mixture was vortexed for 15 mines at room temperature, then diluted by the buffer containing 140 mM KCl and 25 mM HEPES, pH 7.4 for a final OG concentration of 0.33% (w/v). OG was removed by dialyzing the liposome solution in the same buffer using Slide-A-Lyzer™ Dialysis Cassettes (20 kDa cutoff) (Thermo Scientific, 66003) for two days at 4 °C with a buffer change every 16 hours. All SUVs were harvested, stored in 4 °C, and used within three weeks.

**Figure 1.**
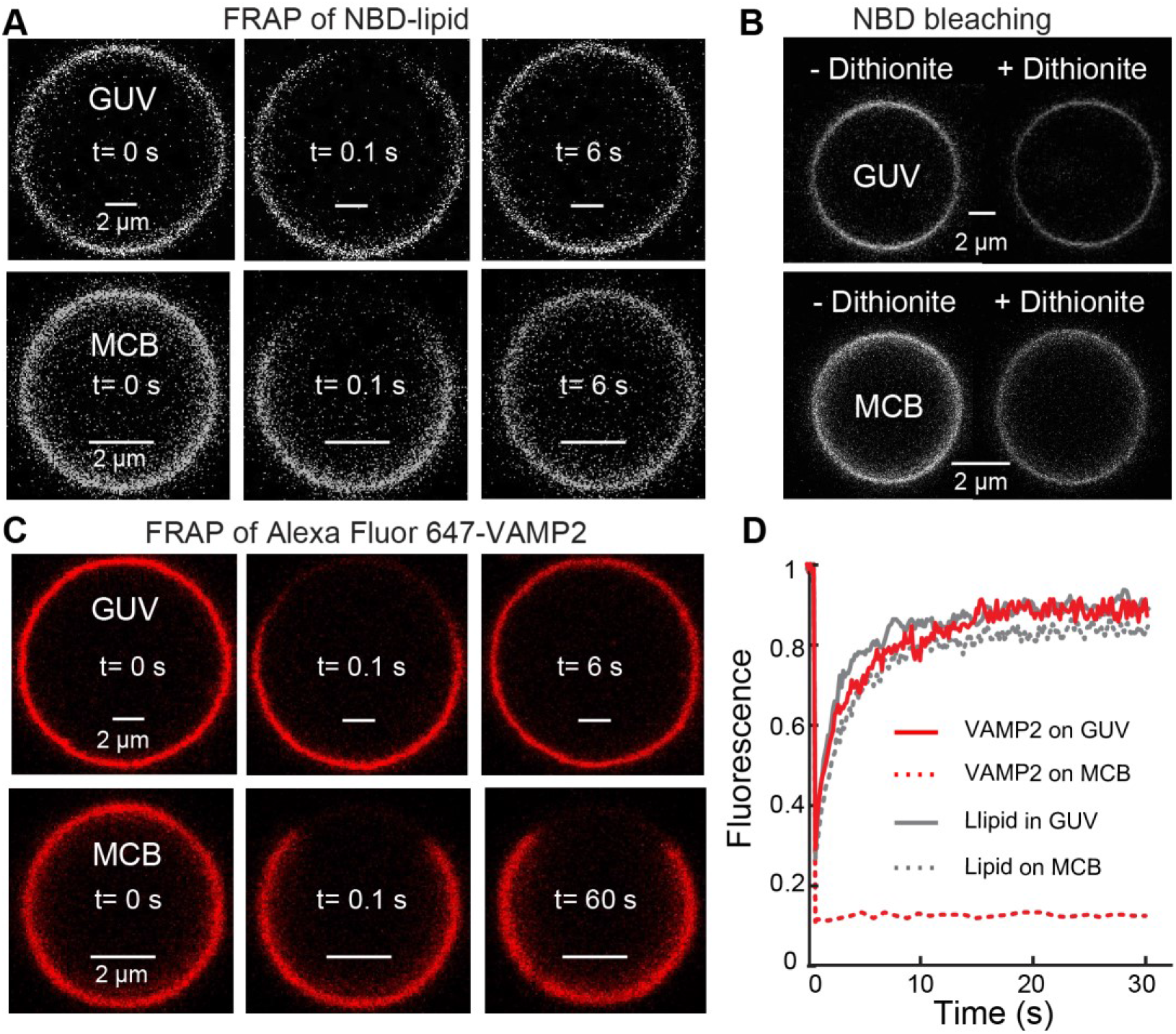
Representative transmembrane protein is immobilized in the bilayer supported on the silica bead and fully mobile in the GUV membrane. (A) Fluorescence images of the lipids in the same GUV or membrane coated on the silica bead (MCB) taken before (t=0) and after photobleaching at the indicated time. (B) Fluorescence images of the lipids in the same GUV or MCB before and after dithionite treatment. (C) Fluorescence images of VAMP2 in the GUV or MCB taken before (t=0) and after photobleaching. (D) Fluorescence intensities as a function of time after photobleaching. The intensities were normalized by the corresponding intensities just before photobleaching.

**Figure 2.**
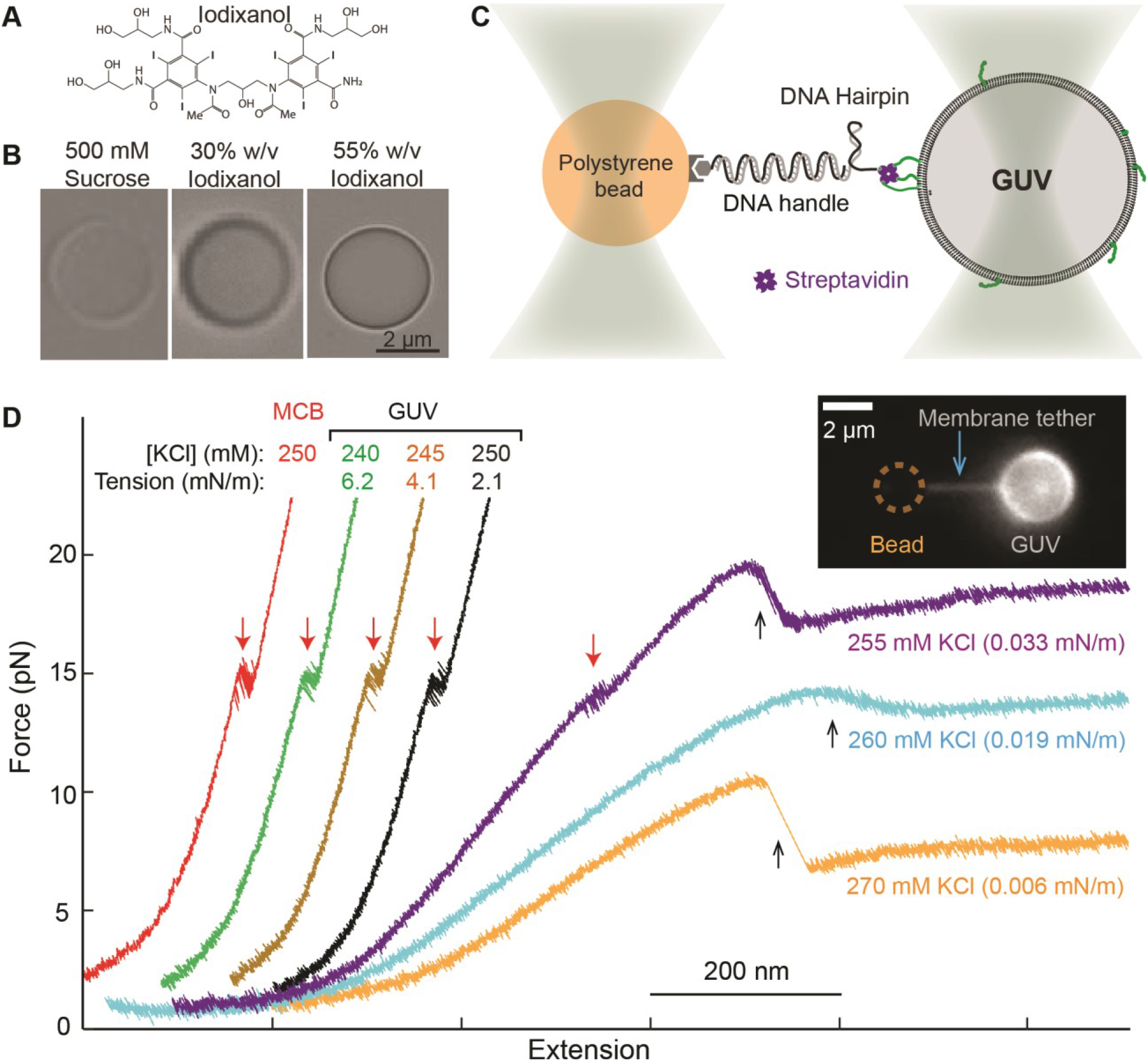
Trapping iodixanol-containing GUVs for single-molecule manipulation. (A) Molecular formula of iodixanol. (B) Bright-field images of optically trapped GUVs containing sucrose only or iodixanol. (C) Schematic diagram of the experimental setup to pull a single DNA hairpin attached to the optical trapped GUV. (D) Force-extension curves (FECs) obtained by pulling the same hairpin DNA attached to the membrane-coated silica bead (MCB) or the same GUV with different membrane tensions in the buffers containing different concentrations of KCl. The three FECs on the left well overlapped the FEC corresponding to 250 mM KCl, but were successively shifted to the left for clarity. Red and black arrows indicate reversible unfolding/refolding transitions of the DNA hairpin and abrupt formation of membrane tethers, respectively. The inset shows the fluorescence image of a membrane tether pulled out of the optically trapped GUV containing Rhodamine-PE.

### Syt1 C2AB and VAMP2 preparation

The sequences and purification of the Syt1 C2AB construct and the full length VAMP2 with single cysteine mutation Q36C were previously described (14, 43). Briefly, the Syt1 C2AB construct contained an Avi-tag at its N-terminus and a unique cysteine at its C-terminus for crosslinking with the DNA handle. The C2AB domain and the thiol-containing DNA handle were crosslinked as previously described (14). VAMP2 and Alexa Fluor 647 maleimide (Thermo Fisher, A20347) was mixed with a molar ratio of 1:3 in the presence of 1 mM tris (2-carboxyethyl) phosphine (TECP) and incubated at room temperature for 1 hour. Then, dithiothreitol (DTT) was added to the mixture to a final concentration of 5 mM to quench unreacted maleimide. Free dye was removed by Micro Bio-Spin 6 Columns (Bio-Rad Laboratories, 7326222).

### Preparation of membrane coated beads

The membrane coated beads (MCBs) were prepared as described elsewhere in detail (14). The MCBs used to pull DNA hairpins contained 99.97 mol% POPC and 0.03 mol% biotin-PE. The VAMP2 SUVs had 99.65 mol% POPC, 0.25% 1,2-dioleoyl-sn-glycero-3-phosphoethanolamine-N-(7-nitro-2-1,3-benzoxadiazol-4-yl) (NBD-PE), and 0.1 mol% Alexa Fluor 647 VAMP2. Briefly, 100 μL of prewashed silica beads (Bangs Laboratories, SS04002 and SS05003) with a diameter of 2.06 μm (for the pulling experiments) or 6 μm (for the imaging experiments) were added into the corresponding 500 μL SUV solution containing 1 mg/ml lipids. SUVs spontaneously bound to and collapsed on the surfaces of silica beads to form supported bilayers. The bead solution was vortexed at 1500 rpm at 37 °C using Thermal Mixer C (Eppendorf) for 1 hour to complete the membrane coating process. MCBs were separated from the excessive liposomes by centrifugating the bead solution at 500 g at room temperature for 1 min to precipitate the beads and then removing the supernatant. The beads were washed three times by adding 1 mL Buffer A, re-suspending the beads, and centrifugation. The MCBs were stored in 100 μL buffer A at 4 °C and used within one week.

### GUV preparation

GUVs containing sucrose only were generated by the electroformation method (33). 20 μL of 99.97 mol% POPC and 0.03 mol% biotin-PE with a final concentration of 5 mg/mL in chloroform was deposited onto platinum electrodes in small drops (~0.5 μL per drop). The lipids were dried in the vacuum desiccator for 1 hour to form lipid films on the electrodes. Then, the electrodes were gently immersed into a plastic tube with a buffer containing either 500 mM sucrose or 1 M sucrose. For the GUVs containing Alexa Fluor 647 VAMP2, 40 μL SUV solution containing 2 mg/ml lipids was deposited onto platinum electrodes in small drops (~0.5 μL per drop). The SUVs solution was first dried in the fume hood for 15 mines and then dried in the vacuum desiccator for 1 hour to form lipid films on the electrodes. An alternating current with a sine wave (function/arbitrary waveform generators, SIGLENT’s SDG2042X) was applied to the platinum electrodes with a peak-to-peak voltage of 2.3 V and frequency of 10 Hz for 4 hours. The GUVs were harvested and stored in 4 °C, which were stable for one week.

The GUVs containing over 30% iodixanol were made by an alternative inverted-emulsion method (44), because of the poor yield of the GUVs generated by the electroformation. A total of 0.4 μmol of lipids containing 84.97 mol% POPC, 10 mol% 1,2-dioleoyl-sn-glycero-3-phospho-L-serine (DOPS), 5 mol% 1,2-dioleoyl-sn-glycero-3-phospho-(1’-myo-inositol-4’,5’-bisphospha-te) (PI(4,5)P_2_), and 0.03 mol% biotin-PE were mixed in chloroform and dried in a clean glass vial. Then 400 μL liquid paraffin was added to the dried lipids and incubated at 50 °C for 1 hour to dissolve the lipids, which yielded a lipid solution in paraffin with a total lipid concentration of 1 mM. 200 μL of the solution was gently deposited on top of 500 μL buffer containing 20 mM Tris, pH7.4, 200 mM KCl, and 55 mM glucose in a 1.5 mL centrifuge tube and incubated at room temperature for 1 hour until the interface between the oil and aqueous phases became flat, where a monolayer of lipids was formed. 20 μL buffer to be encapsulated into the GUVs, i.e., 5 mM HEPES, pH 7.4, 55% (w/v) iodixanol, 210 mM sucrose, was added to the remaining 200 μL lipid solution in paraffin and sonicated for 5 min to prepare the inverted emulsion solution. This emulsion was added on top of the lipid solution in paraffin above the aqueous solution in the centrifuge tube. The mixture was then centrifuged at 1000 g for 5 min to allow water droplets in the emulsion to pass through the lipid monolayer into the bottom aqueous solution to form GUVs. The bottom GUV solution was collected and stored at 4 °C before use. The GUVs used to pull the DNA hairpin were similarly made, which contained 99.97 mol% POPC and 0.03 mol% biotin-PE.

### Estimations of the GUV membrane tension and force constant of deformation

Suppose a GUV has a radius *r* and buffers of osmolarity *C_in_* and *C_out_* inside and outside of the GUV, respectively, with Δ*C* = *C_in_* − *C_out_* > 0. Using van’t Hoff’s law for the osmatic pressure and the Young-Laplace equation for the membrane tension (45), one can derive the equilibrium membrane tension as

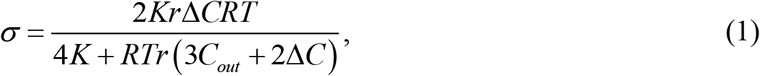

where *K* is the elastic modulus of the GUV membrane (46), *R* the molar gas constant, and *T* the temperature. The membrane tension can also be calculated using the measured equilibrium force of the membrane tether (*f*), i.e.,

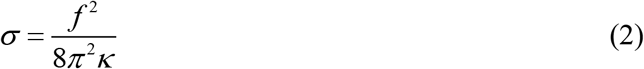

where *κ* is the membrane bending rigidity. The corresponding radius of the membrane tether can be calculated as

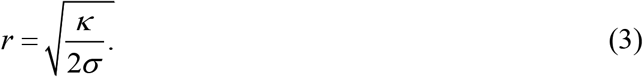

In our estimations, we chose *K* =220 pN/nm, *κ* =10 pN×nm, and *RT* =2.5 kJ/mol.

### Dual-trap high-resolution optical tweezers

The dual-trap optical tweezers were home-built as described elsewhere in detail (47). Briefly, a single laser beam of 1064 nm from a solid-state laser (Spectra-Physics, J20I-BL-106C) was collimated, expanded, and split into two orthogonally polarized laser beams. One of the laser beams was reflected by a mirror attached to a piezoelectrically controlled stage that could turn in two axes, which was used to accurately move the corresponding optical trap in the sample plane. The two beams were further combined, expanded, and finally focused by a water-immersion 60× objective with a numerical aperture of 1.2 (Olympus) to form two optical traps. The outgoing laser beams were collimated by a second identical objective, split, and projected onto two position-sensitive detectors (PSDs, Pacific Silicon Sensor, CA) to detect bead movements through back-focal plane interferometry. The trap stiffness was determined by Brownian motions of the trapped beads. A customized microfluidic chamber with three parallel flow channels was used to deliver beads through the top and bottom channels to the central channel where optical trapping occurred (48).

### Single-molecule experiments

All pulling experiments were performed using the dual-trap high-resolution optical tweezers as previously described (18, 48, 49). Briefly, ~500 ng DNA handles with the biotin labels at the 5’ end, the overhang DNA hairpin, or crosslinked with the Syt1 C2AB domain were mixed with 2 μL 1 mg/mL streptavidin solution with a streptavidin to DNA handle molar ratio of 100:1 in final volume of 5 μL and incubated at room temperature for 15 min. 1~2 μL dilution of such streptavidin-DNA mixture containing 1~10 ng DNA was mixed with 10 μL anti-digoxigenin antibody coated polystyrene beads 2.1 μm in diameter (Spherotech) and incubated at room temperature for 15 min. The beads were diluted in 1 mL buffer A. The stock GUV or MCB solutions from the GUV/MCBs preparation steps were diluted by 10~20 or 1000 fold, respectively, using buffer A with a final volume of 1 mL. Subsequently, the beads, GUV, or MCB solutions were injected into the top and bottom channels in a home-made microfluidic chamber filled with buffer A with oxygen scavenging system containing 55 mM glucose, 0.02 unit/mL glucose oxidase (Sigma-Aldrich), and 0.06 unit/mL catalase (Sigma-Aldrich). For the SUV pulling experiment, 10 μL anti-digoxigenin antibody coated polystyrene beads was mixed with 1 μL 20 ng/μL overhang DNA handle and 9 μL 1 mg/mL SUVs containing oligonucleotide-PE and incubated at room temperature for 15 min. Then, the beads were diluted in 1 mL buffer A and injected into the bottom channel. A single anti-digoxigenin bead from the top channel was trapped and brought close to a single MCBs, GUVs or anti-digoxigenin bead from the bottom channel held in another optical trap to form a single protein (or lipid)-DNA tether between the two beads. The tether was pulled or relaxed by moving one optical trap relative the other fixed trap at a speed of 10 nm/s.

## RESULTS AND DISCUSSION

### Representative integral membrane protein is mobile on GUVs, but not on supported bilayers

We have recently adopted the membrane coated on the surface of the silica bead (MCB) to study membrane binding affinity and kinetics of the C2AB domain in synaptotagmin-1 (Syt1) using optical tweezers (14). In principle, integral membrane proteins can be reconstituted into the supported bilayers and pulled in a direction perpendicular to the membrane surface to study their dynamics as Syt1. However, like in other supported bilayers (27), the integral membrane proteins might suffer from nonspecific interactions with the underlying glass surfaces. This motivated us to examine the lateral mobility of integral membrane proteins in the lipid bilayers coated on silica beads using fluorescence recovery after photobleaching (FRAP). To this end, we chose VAMP2, a SNARE protein of 116 amino acids in length with a single C-terminal transmembrane domain, as a representative for integral membrane proteins (3). We labeled VAMP2 with the Alexa Fluor 647 dye and reconstituted the protein into the bilayer on the surface of a silica bead 6 μm in diameter. For comparison, we also reconstituted the dye-labeled proteins into GUV membranes. Both GUV and supported membranes also contained NBD-labeled lipids. First, we tested the mobility of the lipid in the membranes using FRAP. As expected, after photobleaching NBD in a small region on the GUV or MCB, the fluorescence in the region quickly recovered within ~10 seconds with comparable rates for lipids on both GUV and MCB (Fig. 1A), suggesting that the lipids are fully mobile. Next, we examined the unilamellarity of both membranes. We treated the membrane-coated beads (MCBs) and GUVs with dithionite that specifically quenches the NBD dyes labeled on the lipids in the outer leaflets of the membranes. Comparing bead or GUV images before and after dithionite treatment, we found that their fluorescence intensities decreased by ~50% (Fig. 1B), indicating unilamellar membranes coated on bead surfaces as well as in the GUV. Finally, we tested VAMP2 mobility in the membrane of GUV or on the bead surface using FRAP. The fluorescence of VAMP2 on the GUV in the photobleached region quickly recovered within 10 s (Fig. 1C; Video S1), consistent with the rapid diffusion of VAMP2 proteins in the GUV membrane. In contrast, no fluorescence recovery was observed for VAMP2 on the membrane on the bead surface even 30 minutes after photobleaching (Fig.1, C and D; Video S2). Thus, the VAMP2 proteins were immobilized on the bead surface. Combining with previous results (27–29), our experiments revealed an intrinsic drawback of the bilayers supported on silica beads as a model membrane to study integral membrane proteins using optical tweezers, despite its success in studies of protein-membrane interactions with mobile lipids (14). We thus turned to GUVs and SUVs as potential model membranes to pull membrane proteins.

### Optical tweezers stably trap GUVs containing iodixanol

To trap GUVs for pulling membrane proteins, we planned to increase the GUV trapping strength characterized by the stiffness of the optical trap. Given the size of a micron-sized object and the laser trapping power (typically a few hundred milliwatts), the trap stiffness increases with the refractive index (RI) of the object relative to that of water (RI=1.33) (50). Therefore, we encapsulated solutions with different refractive indices inside GUVs and measured their trapping stiffness based on their Brownian motion in the optical trap with a fixed trapping laser power (51). All GUV membranes contained 99.98 mol% POPC and 0.03 mol% biotin-DOPE. The buffers outside the GUVs contained 20 mM Tris, pH 7.4, 55 mM glucose, and different concentrations of potassium chloride to balance the osmotic pressure of the GUVs. We first tested GUVs encapsulating 0.5 M sucrose (RI=1.36), as they were used in previous trapping experiments (39). We obtained an average trap stiffness of 0.025 ± 0.005 (mean ± SEM) pN/nm for GUVs with diameters of 2.5-3.5 μm (Table 1). The GUV trap was rather weak, compared with the average trap stiffnesses of 0.160 ± 0.006 pN/nm and 0.240 ± 0.006 pN/nm for membrane-coated silica beads (RI=1.45) and polystyrene beads (RI=1.57), respectively, with diameters of ~2 μm. Increasing the sucrose concentration to 1 M (RI=1.38) only slightly enhanced the trap stiffness to 0.045 ± 0.006 pN/nm. Thus, despite being widely used in GUV preparation, sucrose does not significantly enhance GUV trapping due to it low refractive index.

**Table 1.**
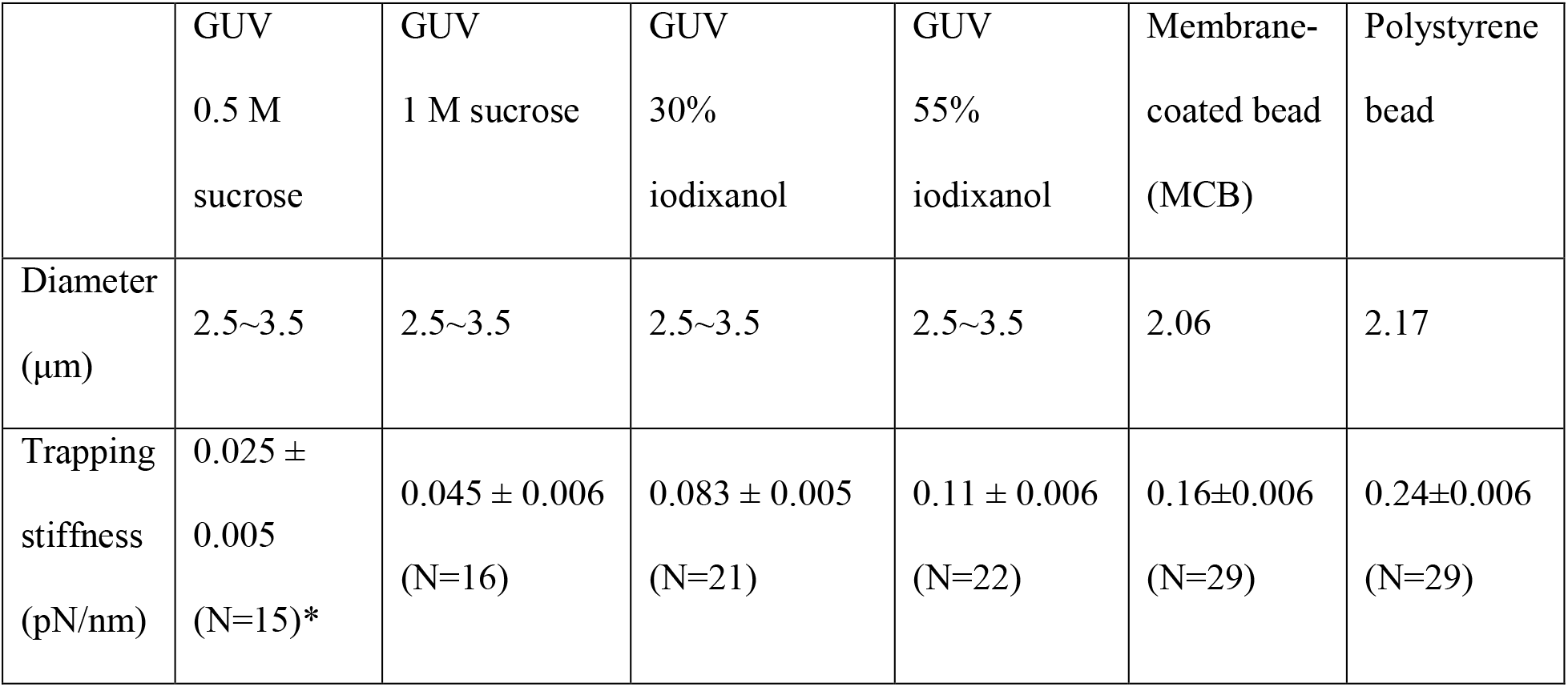
Trapping stiffnesses of GUVs containing sucrose or iodixanol in different concentrations, membrane-coated silica beads, and polystyrene beads. *Number of GUVs or beads tested.

|  |  |  |  |  |  |  |
| --- | --- | --- | --- | --- | --- | --- |
|  | GUV<br>0.5 M<br>sucrose | GUV<br>1 M sucrose | GUV<br>30%<br>iodixanol | GUV<br>55%<br>iodixanol | Membrane-coated bead<br>(MCB) | Polystyrene<br>bead |
| Diameter<br>($\mu\text{m}$) | 2.5~3.5 | 2.5~3.5 | 2.5~3.5 | 2.5~3.5 | 2.06 | 2.17 |
| Trapping<br>stiffness<br>(pN/nm) | $0.025 \pm 0.005$<br>(N=15)* | $0.045 \pm 0.006$<br>(N=16) | $0.083 \pm 0.005$<br>(N=21) | $0.11 \pm 0.006$<br>(N=22) | $0.16 \pm 0.006$<br>(N=29) | $0.24 \pm 0.006$<br>(N=29) |

To promote GUV trapping, we added iodixanol, also known as OptiPrep (Fig. 2A), inside the GUV. The iodixanol solution has widely been used as a medium for density gradient centrifugation and a radiocontrast agent in diagnostic imaging, because of its high density and low osmolarity, viscosity, and toxicity (52, 53). Recently, it has also gained applications in optical imaging due to its high refractive index and low absorbance for visible or infrared light (54). The typical 60% iodixanol stock solution has a high reflective index of 1.43, close to that of silica. The low absorbance is important for GUV trapping, because it minimizes laser heating due to the extremely high laser power density in optical traps (~ 10 MW/cm^2^) (47, 55). We prepared two batches of GUVs, one containing 30% (w/v) iodixanol and 430 mM sucrose and the other, 55% (w/v) iodixanol and 355 mM sucrose. Here sucrose was used to adjust both solutions to approximately equal osmolarity. We made the GUVs by the inverted emulsion method (44, 56), which produced higher yield of GUVs than the alternative electroformation formation method (33). All GUVs could be readily trapped, imaged, and appeared spherical (Fig. 2B). Due to their high refractive index, these GUVs exhibited significant higher contrast than those containing sucrose only. The trapping stiffnesses for GUVs containing 30% and 55% iodixanol were 0.083 ± 0.005 pN/nm and 0.110 ± 0.006 pN/nm, respectively (Table 1). The latter was close to that of membrane-coated silica beads (~0.16 pN/nm), but about half of the stiffness of polystyrene beads (~0.24 pN/nm). Therefore, GUVs containing ≥30% iodixanol could be stably trapped by optical tweezers.

### Pulling single DNA hairpins attached to trapped GUVs

To examine whether the trapped GUVs could further serve as a force and displacement sensor like a bead to directly measure dynamics of macromolecules, we investigated the folding and unfolding dynamics of a DNA hairpin attached to the GUV containing 55% iodixanol. The DNA hairpin contained a stem of 20 bp and a thymidine tetraloop (Fig. 2C). It was directly tethered to the GUV at one end and to the 2.09 μm anti-digoxigenin antibody-coated polystyrene bead at the other end via a 2,260 bp DNA handle. As a force and displacement sensor (57), the GUV needs to be sufficiently rigid to minimize the deformations induced by the pulling force and thermal fluctuations of the membrane. Thus, we controlled the membrane tension of the GUV by changing the concentration of potassium chloride ([KCl]) in the buffer outside the GUV. As [KCl] decreased below 270 mM, both the osmotic pressure and the GUV membrane tension increased in a predictable manner (see Experimental Procedures)(45).

To test the effect of the GUV deformation on the single-molecule manipulation experiment, we pulled the DNA hairpin on the same GUV, but adjusted its membrane tension by varying [KCl] from 240 mM to 270 mM using a microfluidic system (51). The DNA hairpin was being pulled by moving one trap away from the other fixed trap at a speed of 10 nm/sec. At a high membrane tension with low [KCl]s at 240 mM, 245 mM, and 250 mM, the resultant force-extension curves (FECs) were nearly identical, showing clear folding and unfolding transitions of the DNA hairpin at an equilibrium force of ~14.5 pN (Fig. 2D). In addition, all three FECs overlapped the FEC obtained by replacing the GUV with the MCB. Extension trajectories at a constant trap separation or mean force of 14.5 pN also revealed approximately equal extension changes and close folding and unfolding rates (Fig. 3A, top and middle traces). The signal-to-noise ratio (SNR) to detect the hairpin transition on the GUV (4.4) was slightly lower than on the MCB (5.4). These comparisons demonstrate that, at the high membrane tension, the GUV is suited to pulling macromolecules on membranes and detecting their conformational transitions with high resolution. This conclusion implies that the GUV was relatively rigid and minimally deformed in response to a high pulling force. Consistent with this derivation, no significant GUV deformation was observed from the images of GUVs subject to up to 40 pN pulling force (Fig. 3B).

**Figure 3.**
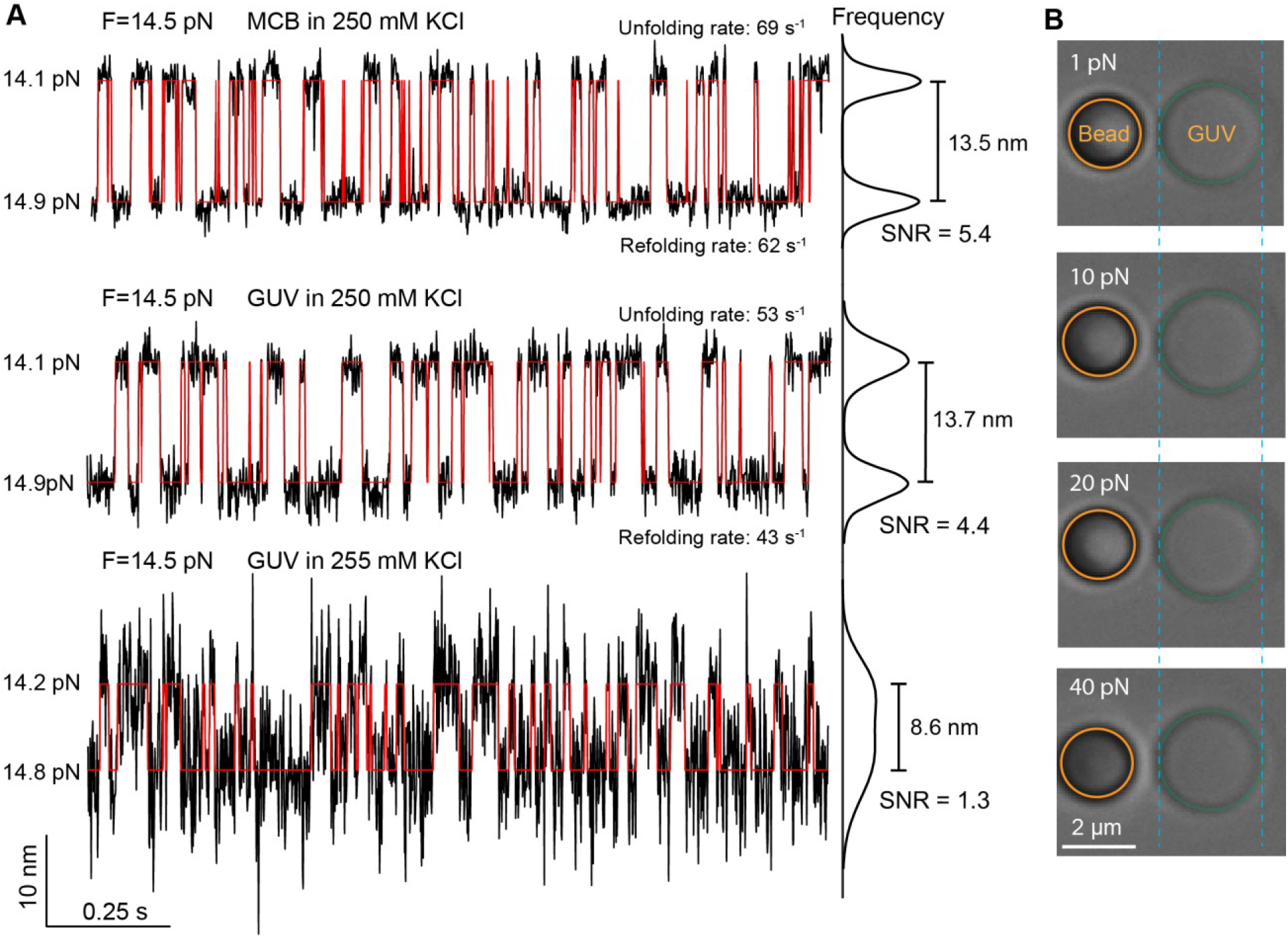
Time-dependent extension trajectories at constant mean force showing reversible unfolding and refolding of the DNA hairpin. (A) Extension trajectories measured when the DNA hairpin was attached to the MCB (top) or GUV in 250 mM (middle) or 255 mM (bottom) KCl concentration. The trajectories were mean-filtered to 1,000 Hz and shown, with their idealized transitions (red lines) derived from hidden-Markov modeling (HMM). The DNA hairpin transition at constant trap separation was accompanied by small force fluctuations. The mean force indicated is the mean of the two average forces corresponding to the folded and unfolded states labeled on the left. On the right are the probability density functions of the extension, which yield the indicated extension changes and the signal-to-noise ratios. (B) Bright-field images of optical trapped GUV and polystyrene bead tethered by the DNA handle and hairpin as in Fig. 2C in the presence of different pulling force.

Theoretical analyses corroborated the negligible GUV deformation induced by the pulling force under our experimental conditions with high GUV membrane tension. Cell or GUV membranes have been used as force probes based on membrane deformation, whose force constant was estimated to be two-fold of the membrane tension (58). Based on the membrane tension measured by membrane tether pulling described in the forthcoming section and the relative [KCl], we derived membrane tensions in the range of 6.2 - 4.1 mN/m for the GUV in 240 - 250 mM KCl, with the corresponding GUV elongation of 1.6-4.8 nm in the presence of 20 pN pulling force (see Experimental Procedures). This contribution to the absolute extension was negligible compared to the ~741 nm extension of the 2,260 bp DNA handle at the same force. For the DNA hairpin transition measured in 250 mM KCl (Fig. 3A), the GUV deformation dampened the extension change by ~0.2 nm, which is significant smaller than the measured 13.7 nm extension change. In conclusion, these calculations corroborated our experimental observations that GUV containing high concentrations of iodixanol with high membrane tensions can be used as a force probe to accurately measure dynamics of macromolecules on membrane surfaces.

### Pulling membrane tethers from trapped GUVs

In contrast, the FECs obtained at lower [KCl] significantly deviated from those described above. At 255 mM KCl, the FEC tilted to higher extension at force below 18 pN (Fig. 2D, purple), indicating significant GUV elongation along the pulling direction, which contributed to the extra extension compared to the extension measured using MCBs at the same force. Although the DNA hairpin transition still equilibrated at ~14.5 pN, the extension change decreased to 8.6, with the corresponding SNR decrease to 1.3 (Fig. 3, bottom trace). Further pulling led to a sudden extension increase and accompanying force decrease (Fig. 2D, purple FEC). Continued pulling only slowly increased force as extension significantly increased. The sudden extension increase and the subsequent approximate force plateau indicate that a membrane tether or nanotubule was being pulled out of the GUV as was confirmed by fluorescence imaging (Fig. 2D, inset). Our observations are consistent with previous experimental results and theoretical analyses based on membrane mechanics (35, 59). As [KCl] was further reduced to 260 mM or 270 mM, the approximate plateau force of the membrane tether decreased with the corresponding decrease in membrane tension, again consistent with previous results (37). Quantitative relationships have been established between the plateau force and the radius of the membrane tether and the membrane tension and bending rigidity (see formulae in Experimental Procedures). Thus, we derived the membrane tensions of the GUV in the three concentrations of potassium chloride and the diameters of the associated membrane tethers (12 nm, 16 nm, and 29 nm at [KCl]s of 255 mM, 260 mM, and 270 mM, respectively). Membrane tethers are widely observed in cells and play important roles in information and material transfer within or between cells (60). They are generated by pulling force and/or various proteins that bind to membranes to sense or generate membrane curvature (36, 61). Thus, the trapped GUVs with low membrane tensions can be used to pull membrane tethers to probe the mechanical properties of membranes or curvature-dependent protein binding and membrane remodeling.

### Synaptotagmin-GUV membrane binding

Next, we asked whether the trapped GUV could be applied to study protein-membrane interactions, using the C2AB domain of Synaptotagmin-1 (Syt1) as our model system. Syt1 C2AB domain binds to membranes in the presence of Ca^2+^ and negatively charged lipids, thereby mediating Ca^2+^-triggered fusion of synaptic vesicles with the pre-synaptic plasma membrane (62, 63). We previously measured the membrane binding energy and kinetics of Syt1 C2AB using MCBs and optical tweezer (14). Therefore, we repeated the experiments by replacing MCBs with GUVs containing 55% iodixanol. As in the previous experiment, the GUVs contained 85 mol% POPC, 10 mol% DOPS, 5 mol% PI(4,5)P_2_, and 0.02 mol% biotin-DOPE. We attached the N-terminus of Syt1 C2AB domain to the GUV membrane through a flexible polypeptide linker and pulled from its C-terminus via the 2,260 bp DNA handle (Fig. 4A). In the presence of 100 μM Ca^2+^ in the solution, the FEC shows reversible membrane binding and unbinding at ~5 pN, followed by sequential unfolding of the C2A and C2B domains at higher force (Fig.4, B and C). The membrane binding was Ca^2+^-dependent, as the signal disappeared when Ca^2+^ was omitted in the solution (Fig. 4B). At constant trap separations, the C2AB binding and unbinding transitions were clearly seen in the force-dependent extension trajectories (Fig. 4D). All these observations are consistent with our previous measurements using MCBs (14), including the equilibrium force and average extension change associated with the C2AB transition. These comparisons indicate that the iodixanol-containing GUVs can be used to study dynamics of membrane proteins in optical tweezers force spectroscopy. Compared with MCBs, the GUVs have several advantages. The transmembrane proteins are fully mobile and free from perturbation by the underlying glass surface. In addition, various macromolecules, small molecules, and buffers can be added to the relatively large interior space of GUVs, which facilitate studies of numerous transmembrane proteins.

**Figure 4.**
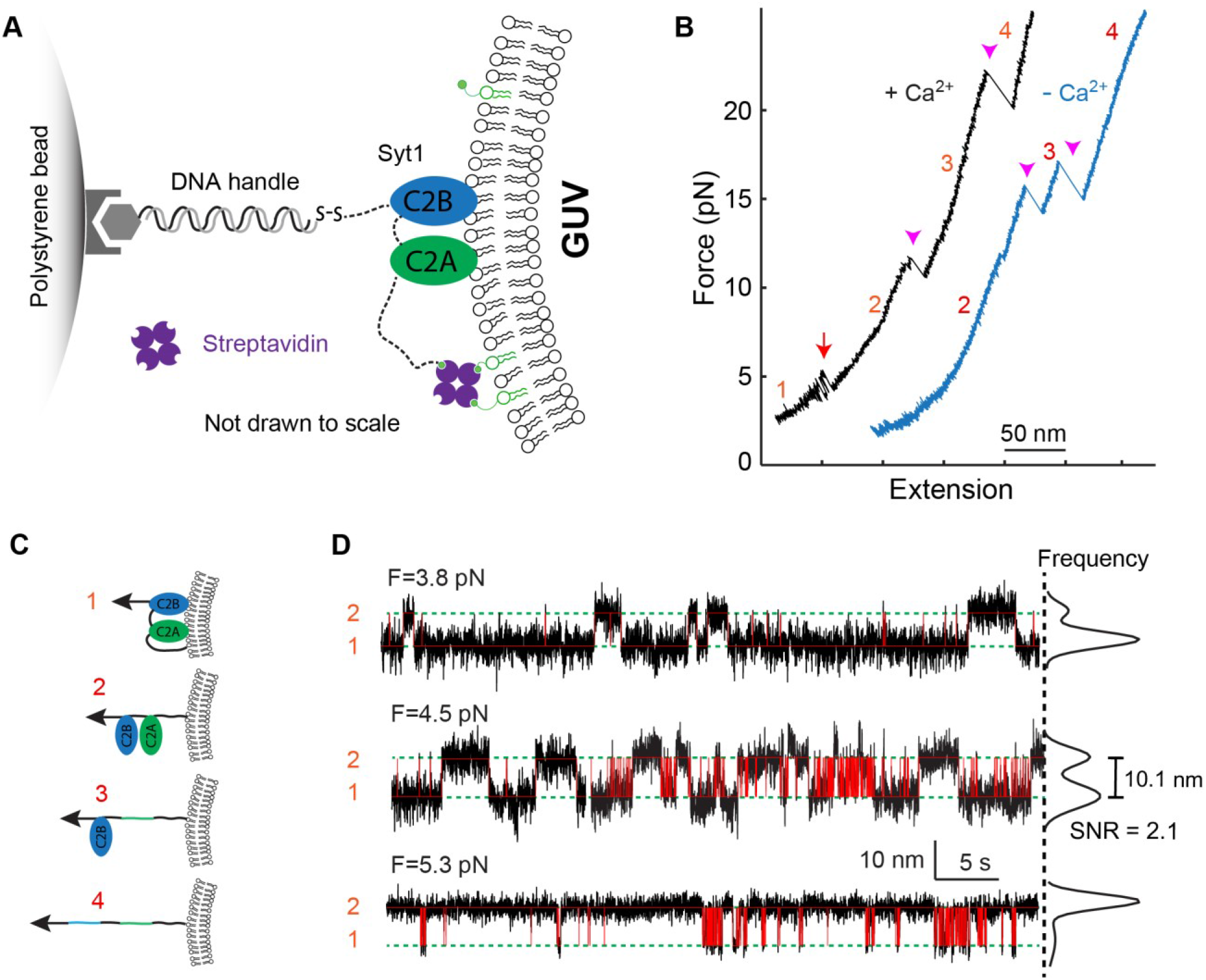
Dynamic membrane binding of Syt1 C2AB domain detected on the surface of the optically trapped GUV. (A) Schematic diagram of the experimental setup. (B) FECs obtained in the presence 100 μM Ca^2+^ (+ Ca^2+^) or absence Ca^2+^ (− Ca^2+^). The red arrow denotes reversible membrane binding and unbinding of Syt1 C2AB domain, and magenta arrow heads indicate unfolding of the C2A and C2B domains. Red numbers label the states associated with different FEC regions as depicted in C. (C) Conformations of different Syt1 C2AB binding or folding states. (D) Time-dependent extension trajectories showing reversible Syt1 C2AB binding to and unbinding from the GUV membrane at constant trap separation or mean force. The trajectories were mean-filtered to 50 Hz and shown with their idealized transitions (red line) derived from HMM.

### SUV as a model membrane to manipulate macromolecules

Despite the advantages of GUVs to manipulate membrane proteins, it is often challenging to reconstitute transmembrane proteins into GUV membranes. In contrast, it is much easier to reconstitute proteins into SUV membranes with well-established protocols (40). In addition, membrane proteins on SUVs can be pulled in directions both perpendicular and parallel to membranes. However, with a diameter in the range of 20-100 nm, SUVs are generally too small to be stably trapped for pulling macromolecules, even with iodixanol encapsulated inside the SUVs. Therefore, we suspended single SUVs between two polystyrene beads using two DNA handles (Fig. 5A). To examine whether potential SUV deformation compromised the single-molecule manipulation experiments, we again pulled the DNA hairpin attached to the SUV surface via one DNA handle. The other DNA handle was directly attached to the SUV lipids through biotin-streptavidin interactions. Wide-field fluorescence imaging of the SUV containing rhodamine-labeled lipids confirmed that a single SUV was being tethered between two beads (Fig. 5B). The FEC of the SUV-DNA tether revealed the characteristic DNA hairpin unfolding and refolding transition at ~14.5 pN that is similar to the transition of the hairpin directly attached to the bead without the SUV (Fig. 5C). The DNA hairpin transition again exhibited an equilibrium force of 14.5 pN (Fig. 5D). A smaller extension change (12 nm vs 13.5 nm, compare with Fig. 2A, top trajectory) and signal-to-noise ratio (3.9 vs 5.4) were expected, because longer DNA handles used here dampened the extension change detected by the beads (64). Therefore, conformational transitions could be accurately measured on the surfaces of SUVs. The observation implied that SUVs are relatively rigid and minimally deform in response to the pulling force. This derivation is consistent with the large force constants of the SUVs in the range of 15-32 pN/nm detected by AFM (65). Using these values, the estimated SUV elongation in the presence of 20 pN was less than 1.3 nm and the extension change of the SUV during the DNA hairpin transition was 0.05 nm. In conclusion, SUVs serve as an excellent model membrane to study dynamics of macromolecules using optical tweezers.

**Figure 5.**
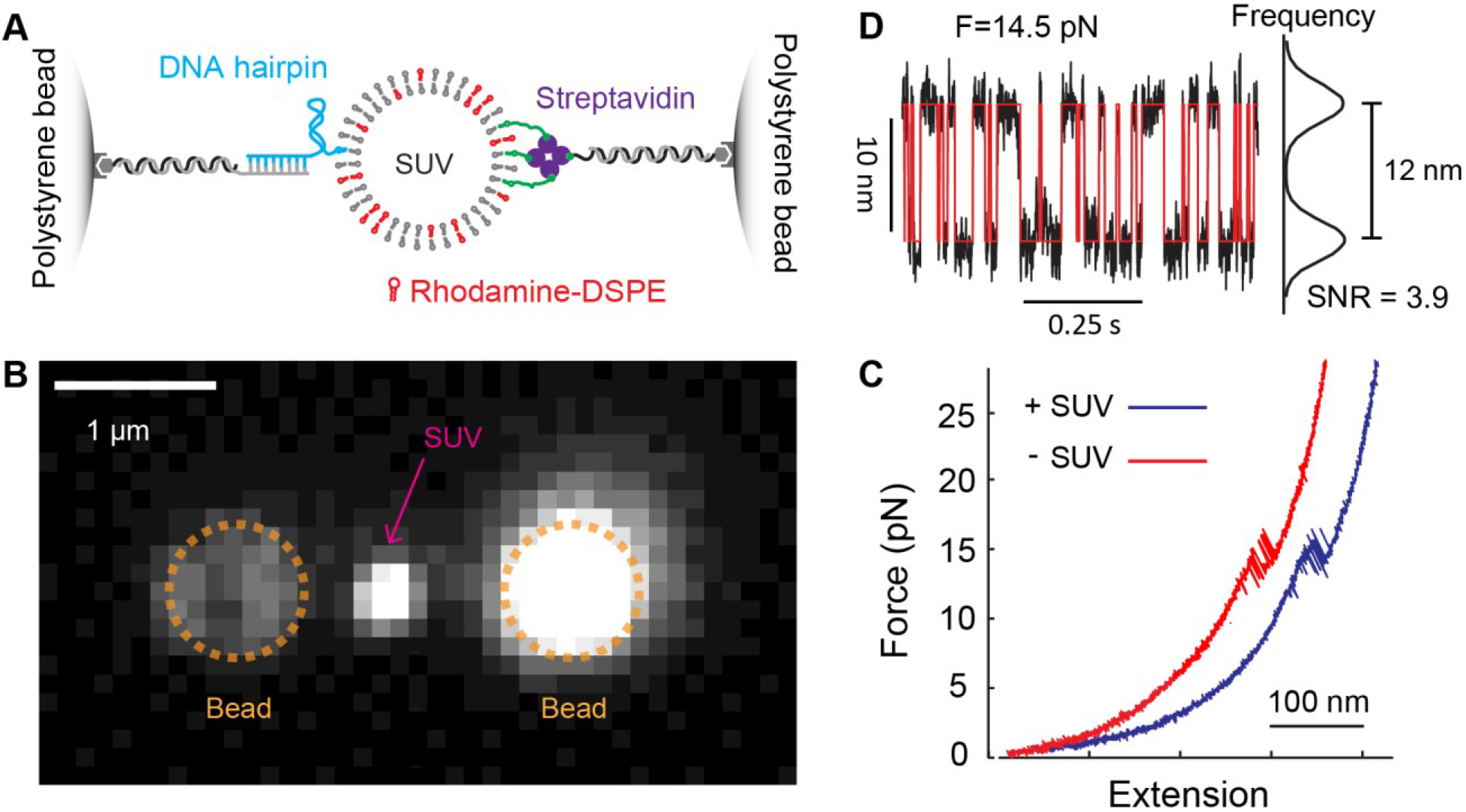
Folding/Unfolding transition of the DNA hairpin detected on the surface of a single SUV tethered between two polystyrene beads. (A) Schematic diagram of the experimental setup to pull the DNA hairpin attached to the tethered SUV. The DNA hairpin was conjugated to the lipid in the SUV. (B) Bright-field fluorescence image of a single rhodamine-labeled SUV tethered between two optical trapped polystyrene beads. Note that SUVs bound specifically to the right bead containing an excess of free biotinylated DNA handles. (C) FEC obtained by pulling the DNA hairpin to high force on the GUV (+ SUV) or directly on the bead (− SUV). (C) Time-dependent extension trajectory at constant mean force showing reversible unfolding/refolding of the DNA hairpin.

## CONCLUSION

Optical tweezers have widely been used to study dynamics of soluble proteins, due to their high resolution and dynamic ranges of measurements for force, extension, and time. In contrast, optical tweezers have rarely been applied to membrane proteins, partly due to lack of model membranes compatible for optical trapping and single-molecule manipulation. We found that iodixanol could be encapsulated inside GUVs to enhance their refractive index, thereby enabling stable trapping of GUVs. The trapped GUVs also served as a model membrane to study dynamics of membranes, proteins, and their interactions. The membrane tension of the trapped GUVs was conveniently regulated by the osmolarity of the buffer outside the GUV, which was facilitated by the microfluidic system used in optical tweezers. We found that GUVs with high membrane tensions were rigid enough to resist significant deformation due to high pulling force, thereby allowing accurate measurements of the extension changes associated with macromolecular conformational transitions around membranes. In a low membrane tension, membrane tethers could be pulled from the trapped GUVs, which could serve as model membranes with tunable curvatures to study curvature-sensitive membrane binding proteins. Membrane tethers have previously been pulled from GUVs aspirated on the tip of micropipettes using optical tweezers (36). However, micromanipulators and other devices are required to control micropipettes. In addition, the GUVs fixed on the sample stage introduce more noise than the optically trapped GUVs due to stage drift (55). Finally, we validated the use of GUVs and SUVs as model membranes in single-molecule manipulation based on optical tweezers with relatively simple model macromolecules, the DNA hairpin and the Syt1 C2AB domain. The methodologies should similarly be applied to more complex membrane proteins, including multi-span transmembrane proteins or protein complexes.

## AUTHOR CONTRIBUTIONS

Y. W., H. J., and Y. Z. designed the experiments, Y. W. and H. J. performed the experiments, Y. W. and Y. Z. analyzed the data, and Y. W., and Y. Z. wrote the paper.

## DECLARATION OF INTERESTS

The authors declare no competing interests.

## SUPPORTING INFORMATION

Supplemental Information includes two videos and can be found with this article online.

## ACKNOWLEDGMENTS

This work is supported by the NIH grant R35 GM131714 to Y.Z. The authors declare no competing financial interests.

## SUPPLEMENTAL INFORMATION

**SUPPLEMENTAL Video 1: FRAP Assay Reveals Free Diffusion of VAMP2 in the GUV Membrane**.

Confocal fluorescence images of the Alexa Fluor 647 labeled VAMP2 in the GUV membrane after photobleaching.

**SUPPLEMENTAL Video 2: FRAP Assay Reveals Immobilized VAMP2 in the Lipid Bilayer Supported on the Silica Bead.**

Confocal fluorescence images of the Alexa Fluor 647 labeled VAMP2 in the lipid bilayer coated on a silica bead 6 μm in diameter after photobleaching.

## Notes

### Competing Interest Statement

The authors have declared no competing interest.

